# Design to Data for mutants of β-glucosidase B from *Paenibacillus polymyxa:* L171M, H178M, M221L, E406W, N160E, F415M

**DOI:** 10.1101/2020.11.17.387829

**Authors:** Xiaoqing Huang, Daniel Kim, Peishan Huang, Ashley Vater, Justin B. Siegel

## Abstract

Computational protein design is growing in popularity as a means to engineer enzymes. Currently, protein design algorithms can predict the stability and function of the enzymes to only a limited degree. Thus, further experimental data is required for training software to more accurately characterize the structure-function relationship of enzymes. To date, the Design2Data (D2D) database holds 129 single point mutations of β-glucosidase B (BglB) characterized by kinetic and thermal stability biophysical parameters. In this study, we introduced six mutants into the BglB database and examined their catalytic activity and thermal stability: L171M, H178M, M221L, E406W, N160E, and F415M.

## INTRODUCTION

Enzyme engineering using computational approaches has had recent success in designing novel proteins not found in nature,^1,2,3^ altering the specificity of protein-ligand interactions,^3^ and improving thermal stability (TS) of proteins.^4,5,6^ However, the predictive ability of current design tools that aim to model catalytic efficacy and even TS are still limited and often rely on human chemical intuition to screen and select designs before testing.

One key challenge in the development of computational design algorithms for improving enzyme function and thermal stability is the lack of a large dataset composed of explicitly measured catalytic parameters such as kinetic constants (*k*_cat_, K_M_, *k*_cat_/K_M_), as well as protein melting temperatures (T_M_). For this reason, we initiated building a database called Design2Data (D2D) to hold kinetic constants and TS of single point mutations of β-glucosidase B (BglB). Previously, Carlin et al. demonstrated using a dataset of 100 kinetically characterized single-point mutants to predict kinetic activity; however, the algorithm was biased towards low activity mutants due to a limited data set.^7^ To minimize this bias and improve the accuracy of functional predictions, the current dataset warranted expansion to improve enzyme engineering tools.

In this study, six new single-point mutations of BglB from *Paenibacillus polymyxa:* L171M, H178M, M221L, E406W, N160E, and F415M were characterized by kinetic activity and thermal stability. Mutations in close proximity to the active site were chosen to examine the impact on substrate binding and structural stability. The molecular modeling tool, Foldit^9^ was used to examine the structure of these variants, as well as to explore the relationship between computational predictions of energetic changes and experimental results of the catalytic and physical properties. In this set of mutants, we hypothesized that we would see a relationship between reduced thermal stability and a decrease in predicted structural stability, as represented by the Total System Energy values calculated from Foldit models. When considering the mutation’s effects on catalytic efficiency, we predicted outcomes ranging from reduced to increased overall catalytic efficiency based on the physiochemical interactions observed on Foldit.

**Figure 1.**
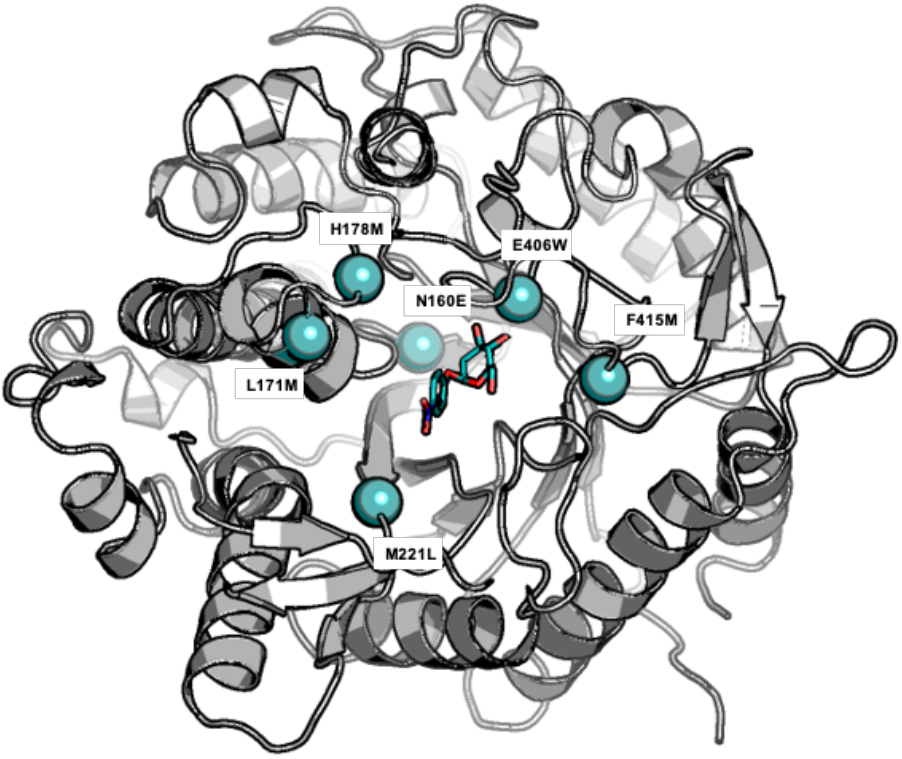
The overview of BglB in complex with reporter substrate 4-nitrophenol β-D glucopyranoside and the locations of six mutants shown as teal spheres. The 3D model structure was generated using PyMOL.^8^

## METHODS

### Designing and modeling of BglB mutants

The six BglB mutants were modeled and chosen using the Foldit Standalone software.^9^ Each mutant design was scored by the Rosetta force-field scorefunction^10^ and was given a Total System Energy (TSE) value. The change in TSE (ΔTSE) relative to the 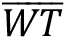 was defined by ΔTSE= TSE (mutant) – TSE 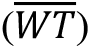. All mutations selected had ΔTSE scores of less than 10, with the exception of E406W (SI Table 1).

### Protein production and purification

Using previously described methods,^11^ the sequence verified BglB mutant plasmids were transformed into chemically competent BLR cells. Following the transformation, the variant expression was induced using isopropyl β-D-1-thiogalactopyranoside (IPTG), lysed, and was purified by metal ion chromatography.^11^ The total yield of the protein was assessed by the A280 BioTek^®^ Epoch spectrophotometer. The purity of the protein samples was analyzed using 4–12% SDS-PAGE (Life Technologies).

### Kinetic efficiency and thermal stability assays

The kinetic assay used the substrate 4-nitrophenyl β-D-glucopyranoside (pNPG) as previously described.^11^ Briefly, the activity was monitored by a spectrophotometer plate reader (Epoch) set to absorbance at 420 nm. The rate of product formation of each mutant was calculated from the Michaelis-Menten model.^12^ The thermal stability assay was performed using QuantaStudio™ 3 System to monitor the fluorescence reading from 30°C to 80 °C. The protein samples were prepared as previously described.^13^ Briefly, the commercially available Protein Thermal shift (PTS)™ kit (Applied BioSystem ®, Thermal Fisher) was used and the experiment was carried by following the standard manufacturer’s protocol. The T_M_ of all mutants were obtained by using the two-state Boltzmann model from the Protein Thermal Shift™ Software 1.3. Pearson correlation coefficient (PCC) analysis was used to examine the relationship between the change in TSE to T_M_ and *k*_cat_/K_M_.^14^

## RESULTS

### Mutants grouping

The six enzyme variants were categorized into two groups (Figure 2A and 2B) based on the distance between the mutation residues and ligand. Group A mutants consisted of L171M, H178M, and M221L, which are further away from the ligand (4Å-11Å). Group B mutants are F415M, N160E, and E406W, which are close to the ligand (<4Å). Foldit modeling predicted variants L171M, M221L, H178M, F415M, and N160E to remain relatively structural unchanged, while E406W resulted in a loss of interactions with the ligand and neighboring residue.

**Figure 2.**
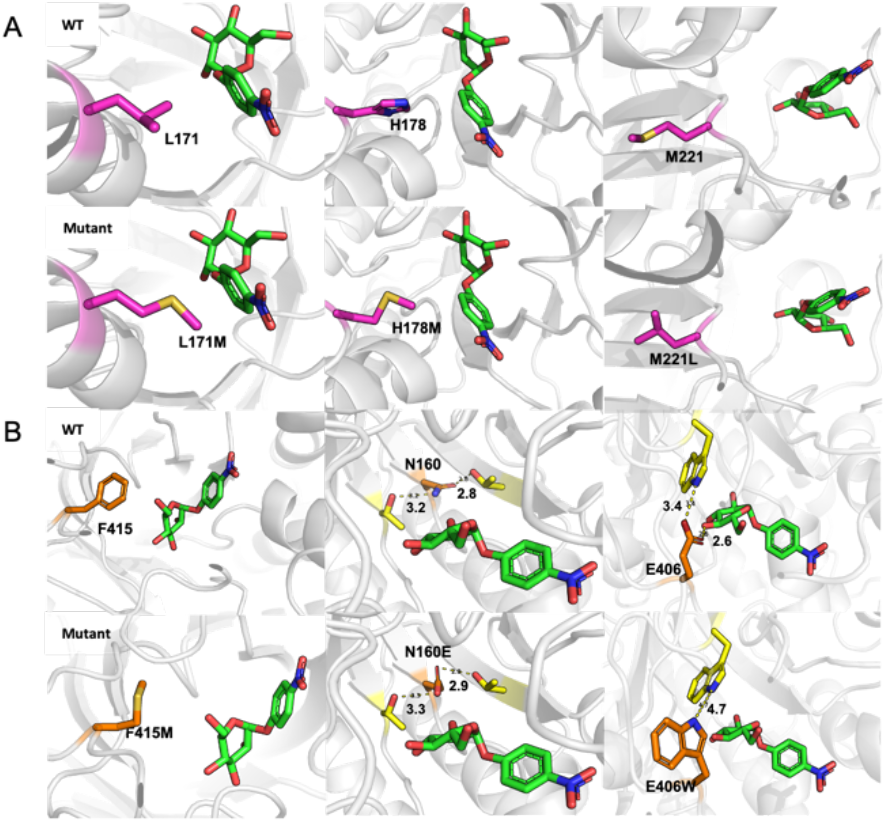
Structural representation of the Wild Types (WTs) and six variants. Ligand was shown in green and the top and bottom panel show WTs and variants, respectively in both sections: A& B. (A) Variant residues: L171M, H178M and M221L shown in pink: (B) Variant residues: F415M, N160E, and E406W shown in orange, with nearby residues shown in yellow. Interaction distances were measured in Å.

### Protein purity and expression

All of the mutants were expressed and purified as a soluble protein. All mutants and WT appear in the SDS-PAGE at 50kD bands which indicate the purity of our samples to being greater than 80% (SI Figure 1).^11^

### Kinetic activity analysis

The catalytic efficiency was measured by the *k_cat_* and K_M_values derived by fitting data to the Michalis-Menten model. The two biological replicates of 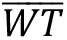 were averaged to be 242.9±187.2 mM^−1^min^−1^ which is consistent with previous experimental values in this system,^7,11^ and considered to be acceptable.^15^ All six mutants showed a decrease in catalytic efficiency compared to the 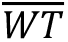 (SI Table 1). The catalytic efficiency for variants L171M, H178M, and M221L decreased within ~2 fold compared to the 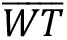, while variants N160E, E406W, and F415M had >100-fold decrease in catalytic activity (Figure 3).

**Figure 3:**
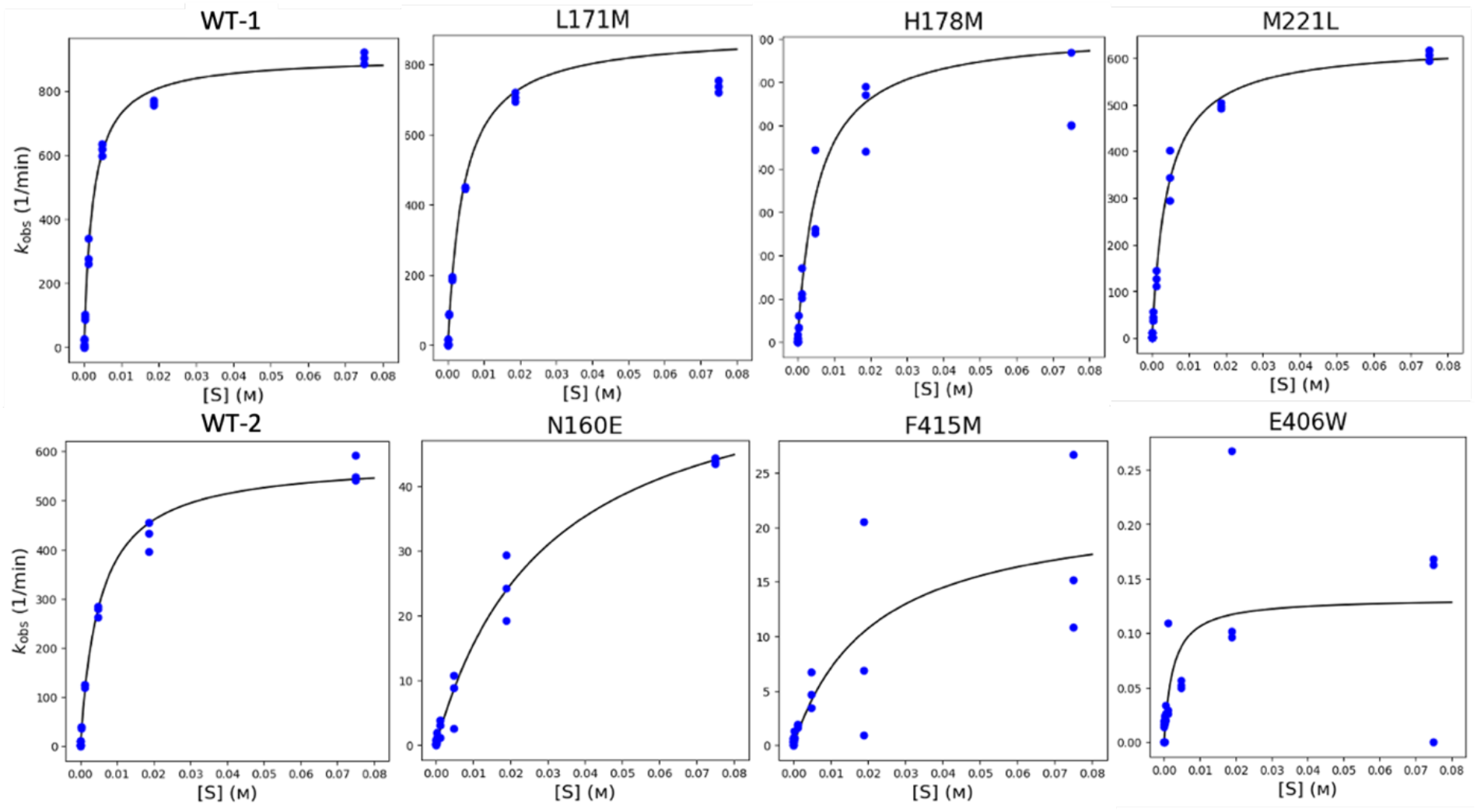
Michaelis-Menten curves of six mutants. Variants L171, H178M, and M221L were tested with WT-1 (top row); N160E, F415M, and E406W were tested with WT-2 (bottom row). The X-axis is the substrate pNPG concentration (M), and the Y-axis represent *k*_obs_ (min^-1^).

### Thermal stability (T_M_) assay

The thermal stability was measured by collecting the triplicated T_M_ values from each mutant. The T_M_ for two biological replicates for 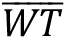 were averaged at 45.7±0.5C (SI Table 1), similar to previous experimental values in this system.^11^ The thermal stability values for L171M, H178M, and M221L are higher than the 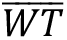 by approximately 1°C. The T_M_ for N160E (49.4±0.5°C) is found higher than the 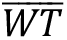 by 3.5 °C. While F415M (44.6±0.2C) and E406W (45.3±0.3°C) are lower than the 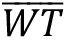 by 1°C and 0.4°C, respectively (Figure 4).

**Figure 4:**
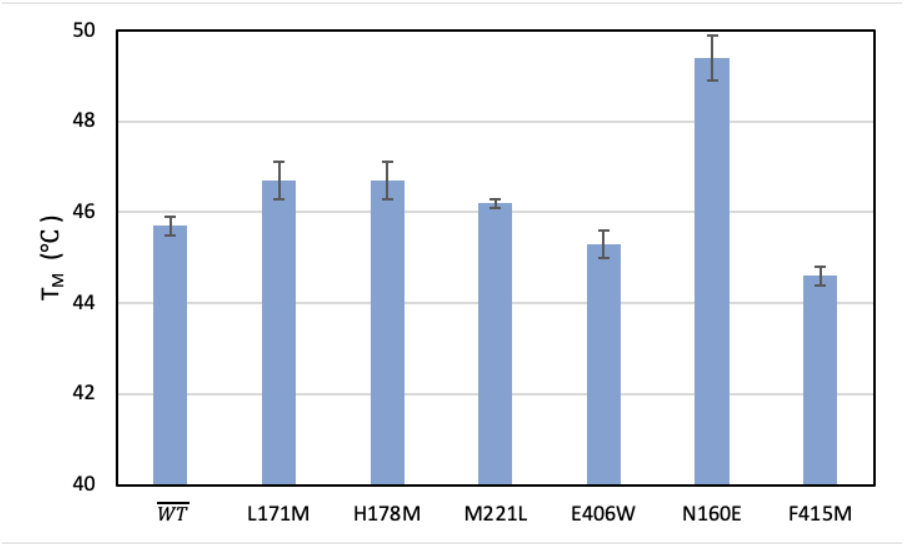
Thermal stability of six mutants and 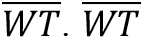 is the average of WT-1 and WT-2. With the exception of N160E, all mutants had around 1°C change in T_M_.

### Relationships between Foldit TSE Score and functional properties

The relationship between ΔTSE and each of the experimental parameters was analyzed using the Pearson correlation coefficient (PCC). The ΔTSE and catalytic activity (Δ*k*_cat_/K_M_) had a PCC of −0.40, while ΔTSE and ΔT_M_had a PCC of −0.89 after removing E406W as shown in Figure 5A and 5B. E406W was removed because ΔTSE (69 points) fell well outside the normal range of the set (0-10 points), suggesting more significant structural changes than were allowed in the simulation.

**Figure 5:**
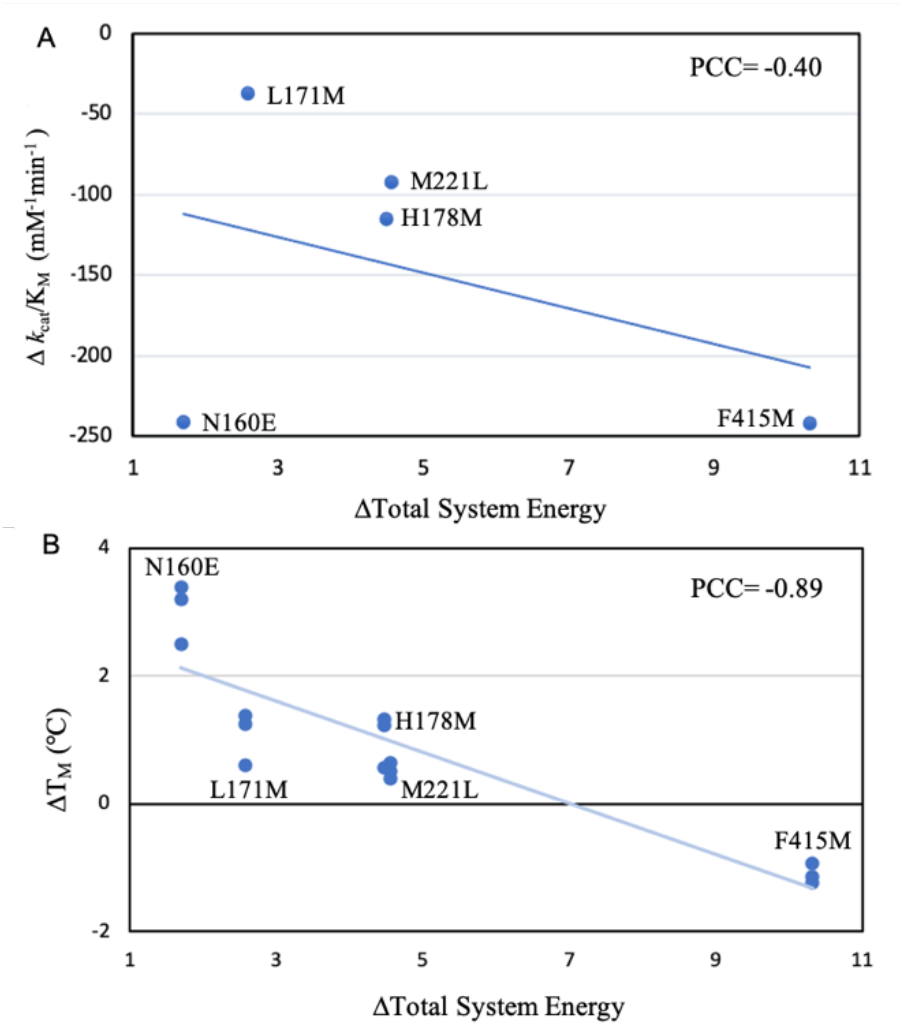
A) The relationship between change in *k*_cat_/K_M_ and change in Total System Energy, data points represent triplicate averages of variants: L171M, H178M, M221L, N160E and F415M. B) The relationship between change in T_M_ and change in Total System Energy, data points show experimental replicates of variants: L171M, H178M, M221L, N160E, and F415M.

## DISCUSSION

When compared with their native forms, L171M and N160E are predicted to have small to modest changes in structural stability (ΔTSE of < 3) while H178M, M221L, F415M, and E406W are predicted to decrease structural stability (ΔTSE of > 4). Two out of six variants (L171M and F415M) are in agreement with this initial hypothesis. Of the mutations that are inconstant with our predictions, N160E has an increase in T_M_ as opposed to the neutral change or modest decrease. Both M221L and H178M had a <1 degree change in T_M_, however previous studies have found that predicted ΔTSE in their range (4.5 and 4.6, respectively) are within a “grey zone range” with low confidence predictions.^13^ Most surprisingly, E406W was predicted to have a ΔTSE ~70 but ended up having a T_M_ of <1 degree. Large predictions, such as this, are most often accurate; however, the algorithms used for those evaluations allowed for more structural sampling than explored in the Foldit simulations.

One of the most prominent relationships we observed in looking at the data in aggregate was the relationship between thermal stability and total system energy change, where we observed a negatively correlative trend (Figure 5B). This was what we expected because we usually associate an increase in thermal stability to an increase in overall protein stability, which was represented by more negative total system energy in Foldit. Another recent BioRxiv publication showed the opposite trend.^16^ Contrasting these two findings exemplifies how small data sets evaluated in isolation like these were susceptible to variations that might be misleading and supporting the need for deeper investigation.

In a closer examination, we observed three notable changes in intermolecular interaction and structural changes between the native and variant enzymes. Mutant F415M was not expected to affect catalytic activity because there were no observed changes in hydrogen bonding between neighboring residues and the ligand. When taking a closer look through PyMOL and Foldit, the change of phenylalanine to methionine seemed to have caused steric crashing with another methionine residue nearby at position 323. This steric crashing could’ve led to structural changes at pocket near the active site causing a large reduction in catalytic efficiency.

Mutant N160E stood out since, unlike the other variants, significant changes in hydrogen bonding to multiple residues in the active site occurred. Specifically, there is predicted to be an unfavorable change in hydrogen bonding angle with nearby residues such as T218 and T116. Somewhat remarkably, variant N160E’s melting point (49°C) fell outside the range of the set (44.6-46.7°C). So, while the mutation had a nearly catastrophic effect on catalytic efficiency it did appear to increase stability. As noted earlier, allowing for more structural flexibility may reveal alternative protein fold stabilizing conformations, which may be possible, but it could prevent the enzyme from folding into a catalytically competent state.

Mutant E406W was hypothesized to have a considerable loss in both catalytic efficiency and thermal stability, simply based on its highly unfavorable predicted change in energy. However, the experimental results did not fully agree with our prediction. While the catalytic activity of the enzyme was nearly extinguished, the T_M_ remained similar to the wild type average. This was not expected because the large increase in the Foldit TSE score suggested a considerable reduction in the enzyme stability to the point of possible unfolding. Structural analysis of this mutant showed the mutant tryptophan might have played a role in causing large steric clashes with the ligand, producing an overall high ΔTSE and blocking ligand binding. Comparing the apo and holo structures could reveal the mutation being compatible with protein structure but not a catalytically competent binding orientation of the substrate.

In conclusion, these six mutants help the development of building towards a larger dataset for benchmarking new algorithms and using molecular modeling tools for understanding the mutational effect on functions. For further improvements, when modeling mutations on residues closer to the active site, the protein should be modeled in both apo and holo states to provide better predictability on stability. Analysis of this small dataset suggests limited predictive capacities of the modeling software we used, thus, pointing to a need for more data to refine the algorithm as well as the need to explore additional simulation protocols.

## AUTHOR INFORMATION

### Author Contributions

Conceived and designed experiments: XH DK PH AV JBS. Performed the experiments: XH DK.

Supervision: PH AV JBS.

Writing - original draft: XH DK PH AV Writing - review& editing: XH DK PH AV JBS.

## ACKNOWLEDGEMENTS

This work was supported by the University of California Davis, the National Institutes of Health (R01 GM 076324-11), the National Science Foundation (award nos. 1827246, 1805510. and 1627539), and the National Institute of Environmental Health Sciences of the National Institutes of Health (award no. P42ES004699). The content is solely the responsibility of the authors and does not necessarily represent the official views of the National Institutes of Health, National Institute of Environmental Health Sciences, National Science Foundation, or UC Davis

## Supplementary Information

**SI Table 1:** Total system energy (TSE), kinetic parameters, and thermal stability (T_M_) of two WTs and six mutants. 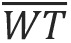 represents the average values of two WTs. The standard errors are the standard deviation of three independent measurements.

| Protein | TSE | ΔTSE | Expression<br>(mg/mL) | $K_M$ (mM) | $k_{cat}$(min<sup>-1</sup>) | $k_{cat}/K_M$<br>(mM<sup>-1</sup>min<sup>-1</sup>) | $T_M$ (°C) |
| --- | --- | --- | --- | --- | --- | --- | --- |
| $\overline{WT}$ | -1089.7 | 0 | 2.1±0.4 | 3.9±0.3 | 744±9.5 | 242.9±187.2 | 45.7±0.5 |
| WT-1 | -1089.7 | 0 | 1.8 | 2.4 | 906.4 | 375.2 | 45.8±0.7 |
| WT-2 | -1089.7 | 0 | 2.4 | 5.3 ± 0.4 | 581.6 ± 13.5 | 110.5 | 45.5±0.1 |
| N160E | -1088.0 | 1.7 | 0.9 | 21.4 ± 52.4 | 61.4 ± 29.0 | 2.1 | 49.4±0.5 |
| L171M | -1087.1 | 2.6 | 2.4 | 4.3±0.3 | 887.4±24.3 | 205.7 | 46.7±0.4 |
| H178M | -1085.2 | 4.5 | 1.3 | 5.6±1.9 | 719.0±108.4 | 128 | 46.7±0.4 |
| M221L | -1085.1 | 4.6 | 4.8 | 4.2 | 630.4 | 150.6 | 46.2±0.1 |
| F415M | -1079.4 | 10.3 | 0.9 | 29.6 ± 20.5 | 22.2 ± 34.3 | 1.0 | 44.6±0.2 |
| E406W | -1020.9 | 68.8 | 1.4 | 2.5 ± 3.1 | 0.1 ± 0.1 | 0.1 | 45.3±0.3 |

**SI Figure 1.**
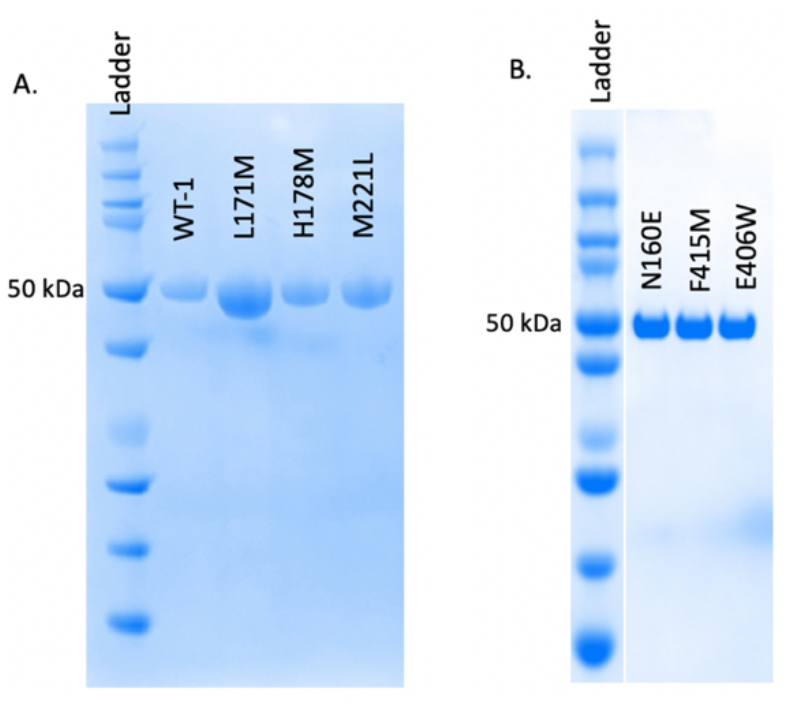
A) SDS-PAGE of L171M, H178M, M221L and WT-1. B) SDS-PAGE of N160E, F415M and E406W.

